# A novel uN2CpolyG Transgenic Mouse Model Recapitulates Multisystemic polyG Proteinopathy Pathology of Neuronal Intranuclear Inclusion Disease

**DOI:** 10.64898/2026.02.10.705201

**Authors:** Yalan Wan, Yilei Zheng, Chao Gao, Yuxuan Lu, Fuze Zheng, Zhen Yu, Jiaxin Wang, Biyu Yang, Jie Zheng, Yun Yuan, Daojun Hong, Nicolas Charlet-Berguerand, Jiaxi Yu, Zhaoxia Wang, Jianwen Deng

**Affiliations:** Department of Neurology, Peking University First Hospital, 100034, Beijing, China; Neuroscience Research Institute and Department of Neurobiology, School of Basic Medical Sciences, Peking University, Beijing, China; Key Laboratory for Neuroscience, Ministry of Education/National Health Commission, Peking University, Beijing, 100083, China; Rare Diseases Medical Center, Peking University First Hospital, 100034, Beijing, China; Department of Neurology, The First Affiliated Hospital of Nanchang University, Nanchang, 330006, China; Institut de Génétique et de Biologie Moléculaire et Cellulaire (IGBMC), INSERM U 1258, CNRS UMR 7104, University of Strasbourg, F-67404 Illkirch, France; Beijing Key Laboratory of Neurovascular Disease Discovery, Beijing, 100034, China

**Keywords:** Neuronal intranuclear inclusion disease, CGG repeat expansion, polyglycine diseases

## Abstract

Neuronal intranuclear inclusion disease (NIID) is a polyglycine disease that primarily affects the neuronal and neuromuscular systems. Here, we developed a novel transgenic mouse model that faithfully recapitulates the multisystemic impairments associated with polyG intranuclear inclusions. Our findings demonstrate that polyG expression induces neurodegeneration, behavioral deficits, and age-dependent accumulation of uN2CpolyG aggregates across multiple tissues.

## Introduction

PolyG diseases represent a newly recognized group of neuromuscular and neurodegenerative disorders, characterized by the accumulation of toxic polyglycine-containing proteins in the nucleus [1]. These diseases spectrum encompasses fragile X-associated tremor/ataxia syndrome (FXTAS) [2], neuronal intranuclear inclusion disease (NIID) [3-6], oculopharyngodistal myopathy (OPDM) [5, 7-12] and spinocerebellar ataxia type 4 (SCA4) [13, 14]. They share a unifying molecular pathogenesis: pathological expansions of CGG/CCG trinucleotide repeats located either in small ORFS hidden within the 5’ untranslated regions (UTRs) of the *FMR1*[15], *NOTCH2NLC* [3-6] and *GIPC1* genes [7, 12], or in the coding region of *ZFHX3* [13, 14] . These expanded CGG repeats are translated into novel and diverse polyglycine-containing proteins that progressively form insoluble inclusions in the nucleus [1, 2, 13, 14, 16-21]. Substantial evidence from cellular, Drosophila, and mouse models have demonstrated the neurotoxic potential of these polyG aggregates. Ubiquitin- and p62-positive intranuclear inclusions, primarily composed of polyG proteins, represent the key pathological hallmark in polyG diseases. Nevertheless, the underlying pathogenic mechanisms associated with these polyG intranuclear inclusions remain largely elusive.

The CGG repeat expansion in the *NOTCH2NLC* gene was recently identified as the genetic cause for NIID/OPDM3, and it represents one of the most common genetic causes of polyG diseases in East Asia [3-5, 22-24]. Notably, in NIID/OPDM3, pathological protein deposition exhibits systemic distribution, occurring not only in neuronal and myofiber nuclei but also in visceral organs, correlating with diverse non-neurological manifestations. These pathological proteins originate from the translation of expanded CGG repeats in the **<u>u</u>**pstream open reading frame of ***<u>N</u>****OTCH****<u>2</u>****NL****<u>C</u>*** into a polyglycine-containing protein named uN2CpolyG, which forms intranuclear inclusions in multiple systems [17, 18]. However, a transgenic mouse model that continuously and generationally expresses toxic uN2CpolyG protein across multiple systems, thereby recapitulating the multisystemic involvement and extensive uN2CpolyG aggregates observed in various organs of NIID, has yet to be established.

In this study, we established a transgenic mouse model which continuously and generationally expressing uN2CpolyG proteins across multiple systems. The transgenic mice exhibit ubiquitous polyG pathology, neurodegeneration and behavioral deficits reminiscent of NIID / OPDM3.

## Meterials and Methods

### Plasmid constructs

The upstream open reading frames (uORFs) of human *NOTCH2NLC*, specifically uN2C and uN2CpolyG, which were reported in the previous studies [17, 25], were subcloned into either the PiggyBAC transposon vector system or the pADM-CMV-C-FH-mCMV-copGFP vector. The uN2C variant containing a 9-glycine repeat was designated as the wild-type uN2C, while the uN2CpolyG variant harboring a 100-glycine repeat was designated as the mutant form, referred as uN2CpolyG. Transgenic mice expressing either uN2C or uN2CpolyG were generated utilizing the PiggyBAC transposon vector system. Additionally, the adenovirus AdMax system (Vigenebio), engineered to express either uN2C or uN2CpolyG, was employed for transduction experiments following the manufacturer’s protocols.

### Transgenic mice construction

The transgenic mice expressing either uN2C or uN2CpolyG were generated by a PiggyBAC transposase system in a C57BL/6J mouse background. Specifically, a microinjection expression plasmid was constructed as follows: CAG (promoter) –uN2C or uN2CpolyG (expression element) - WPRE (mRNA stabilization element) - polyA (termination sequence). PiggyBAC mRNA was synthesized through in vitro transcription. Subsequently, the PiggyBAC mRNA and the vector plasmid harboring either uN2C or uN2CpolyG were co-microinjected into fertilized C57BL/6J mouse oocytes. This process facilitated the random integration of the plasmid into the genome of the C57BL/6J mice, giving rise to transgenic Founder (F0) mice. The microinjection procedure was carried out by Shanghai Model Organisms Center Inc. For genotyping, genomic DNA was extracted from mouse tail biopsies, and PCR analysis was performed to identify transgenic animals. The upstream primer sequence used was GATGGCTATGAACCCTGTGG, while the downstream primer was GCACGGGGGAGGGGCAAACAACAG. The positive band was 716 bp in size, and sequencing of the PCR products confirmed PCR-positive founder (F0) mice expressing uN2C or uN2CpolyG. Expression levels were quantified using RT-qPCR, the upstream primer sequence used was GATGGCTATGAACCCTGTGG, while the downstream primer was AAGCAGCGTATCCACATAGC. Due to marked differences in expression levels (Figure 1b) and the observation that mice exhibiting high expression of uN2CpolyG died before reaching maturity, we selected a uN2CpolyG transgenic line with moderate expression levels—specifically, line uN2CpolyG-3—alongside a uN2C control line (uN2C-2) with comparable expression levels for further analysis. These two mice were then crossed with C57BL/6J wild type mice to establish transgenic lines up to the F5 generation, ultimately generating mice that overexpress uN2C and uN2CpolyG.

**Figure 1.**
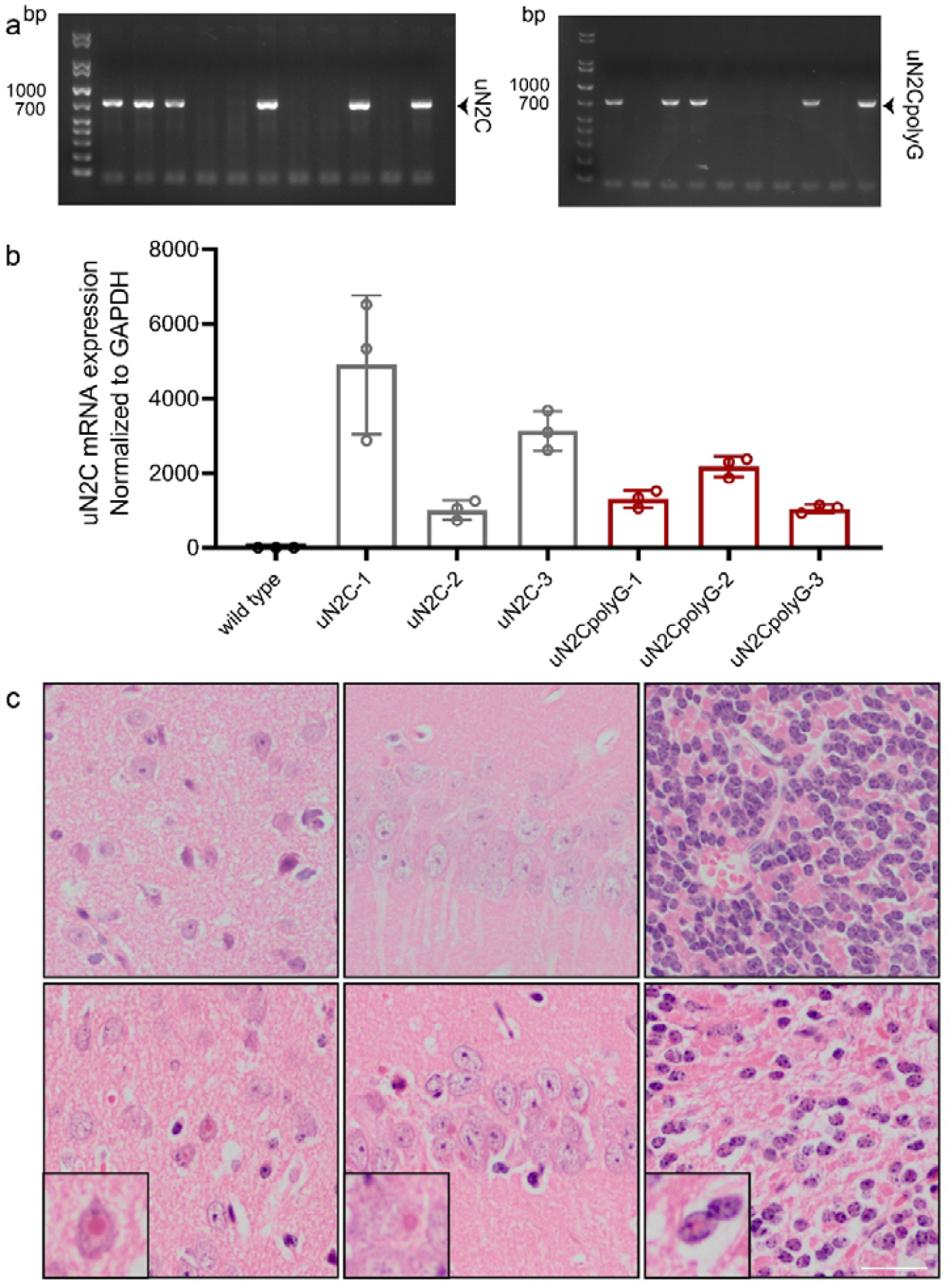
Molecular and neuropathological characterization of uN2C and uN2CpolyG transgenic mice. (a) Genotyping of uN2C and uN2CpolyG mice using genomic DNA from mouse tails by PCR. (b) The relative expression levels of transgene for the uN2C and uN2CpolyG lines. (c) Different brain regions from uN2C and uN2CpolyG mice with hematoxylin and eosin (H&E) staining. Scale bar, 50 μm.

### Immunofluorescent staining and Nissl staining

Mice were anesthetized and perfused intracardially with 0.9% saline solution, followed by 4% paraformaldehyde in 0.1 M phosphate buffer at pH 7.2. Isolated mice brains were dehydrated in 30% sucrose at 4°C and sectioned at 10 μm for subsequent immunofluorescence study: Brain sections were blocked in 10% normal goat serum for 60 min, and then incubated overnight at 4°C with the following primary antibodies: anti p62 antibody (Proteintech, 18420-1-AP, 1:200), anti GFAP antibody (Abcam, ab68428, 1:200), anti IBA-1 antibody (Proteintech, 10904-1-AP, 1:200), 4D12 antibody[25] (specific to uN2CpolyG, 1:100). After incubation with the fluorescent secondary antibodies and 4′,6-diamidino-2-phenylindole (DAPI), the brain sections were mounted to coated glass slides and examined using an inverted research microscope ECLIPSE Ti2-E (Nikon) or a Nikon A1MP confocal microscope.

### Electron microscopy

Electron microscopy analysis was performed as described previously [25]. Briefly, samples were fixed in 2.5% glutaraldehyde overnight at 4°C. Following fixation, the samples were washed with PBS and subsequently fixed in 1% osmium tetroxide (OsO□) for 2 hours. They were then dehydrated through a graded series of ethanol solutions and embedded in Epon812 resin (SPI). Ultrathin sections (70 nm) were obtained using a Leica EM UC6/FC6 ultramicrotome, mounted onto carbon/formvar-coated copper grids, and double-stained with uranyl acetate and lead citrate. Finally, imaging was performed using a Tecnai™ Spirit transmission electron microscope (FEI).

### Mice behaviour analysis

The studies involving animals were reviewed and approved by the Institutional Animal Care and Use Committee (IACUC) of Peking University Health Science Center (BCJE0146). We performed blinded behavioral assessments on 2-month-old male mice expressing uN2C, uN2CpolyG, or wild-type controls (n ≥ 7 per genotype) over two consecutive weeks at the Peking University Health Science Center. During the first week, grip strength testing was conducted on day 1, followed by rotarod performance assessment on day 3. In the second week, open-field testing was carried out on day 1, and Y-maze test was performed on day 3. Prior to each test, all mice were acclimated for 1 hour and subsequently returned to their home cages. The same testing protocol was repeated in 4-month-old male mice. Throughout the study, animals were monitored regularly at high frequency (typically every 2-3 days) for body weight and survival, with additional assessments triggered by clinical signs. Mice were randomly assigned to their respective treatment groups, and data collection was performed in a blinded manner.

### Grip strength tests

The grip force was measured using a digital grip strength meter (SA417, Sansbio, China). Mice were placed on the grip strength meter induction crossbar. After the animals grasped the crossbar, they were dragged horizontally backwards with the tail until their limbs loosened. The average value was recorded as the absolute grip of mice limbs.

### Rotarod tests

In the rotarod test, mice underwent training for 10 minutes per trial with three trials conducted daily over three consecutive days. The rotational speed was progressively increased: 5 rpm on day 1, 10 rpm on day 2, and 20 rpm on day 3. On the testing day, a fixed-speed protocol at 25 rpm was administered three times for each mouse, with each trial lasting up to 10 minutes and an inter-trial interval of 10 minutes. The latency to fall was recorded for each trial.

### Open Field Tests

Each mouse was individually placed in an open-field arena (40 cm × 40 cm × 40 cm) illuminated at 60 lux, with its behavior videotaped for 10 minutes. The following parameters were quantified using specialized software compatible with the behavioral testing chamber (Ji Liang Biotech, China): time spent and distance traveled in the central area (20 cm × 20 cm), as well as total time and total distance traveled throughout the entire arena. The testing arena was cleaned with 75% ethanol between trials to eliminate olfactory cues.

### Y-maze test

Short-term spatial memory was evaluated using spontaneous alternation behavior in a Y-maze, a well-established measure of spatial working memory[26]. Briefly, each mouse was placed at the center of the maze and allowed to explore freely for 5 minutes. An arm entry was recorded only when the mouse’s hind paws were fully within an arm. The sequence and total number of arm entries were documented. An alternation was defined as three consecutive entries into three different arms. Following this logic, the maximum number of possible alternations was calculated as the total number of arm entries minus two. The percentage of spontaneous alternation was then determined as (actual alternations / maximum alternations) × 100%.

### Statistics

Data were shown as mean ± SD for the number of observations. Comparisons of two groups were performed using Student’s unpaired t test for repeated measurements to determine the levels of significance for each group. The one-way ANOVA test is used to calculate the degree of significance wherever there were more than two groups (Dunnett’s multiple comparison test). Each experiment was repeated at least three times independently, and probability values of P < 0.05 were considered statistically significant.

## Results

### Expression of uN2CpolyG causes progressive neurodegeneration in transgenic mice

The *NOTCH2NLC* CGG repeat microsatellite sequence is located within a short upstream open reading frame (uORF) initiated by a classical AUG start codon. Under normal conditions, this sequence contains nine CGG repeats (as annotated in RefSeq) and encodes a small protein of approximately 6 kDa, designated as uN2C. However, in individuals with NIID/ OPDM3, the CGG repeat tract expands to between 60 and 300-500 repeats, leading to the production of an elongated protein. In this expanded form, each GGC codon encodes a glycine residue, resulting in a protein termed uN2CpolyG [25]. Ubiquitin- and p62-positive intranuclear inclusions of uN2CpolyG proteins are observed in multiple systems, representing a key pathological hallmark of this disease [17, 18]. To investigate the pathogenic role of uN2CpolyG intranuclear inclusions, we developed a humanized transgenic mouse model according to previous studies [17, 25] (Figure 2a). Genotypes were confirmed by polymerase chain reaction (PCR) and DNA agarose gel (Figure 1a).

**Figure 2.**
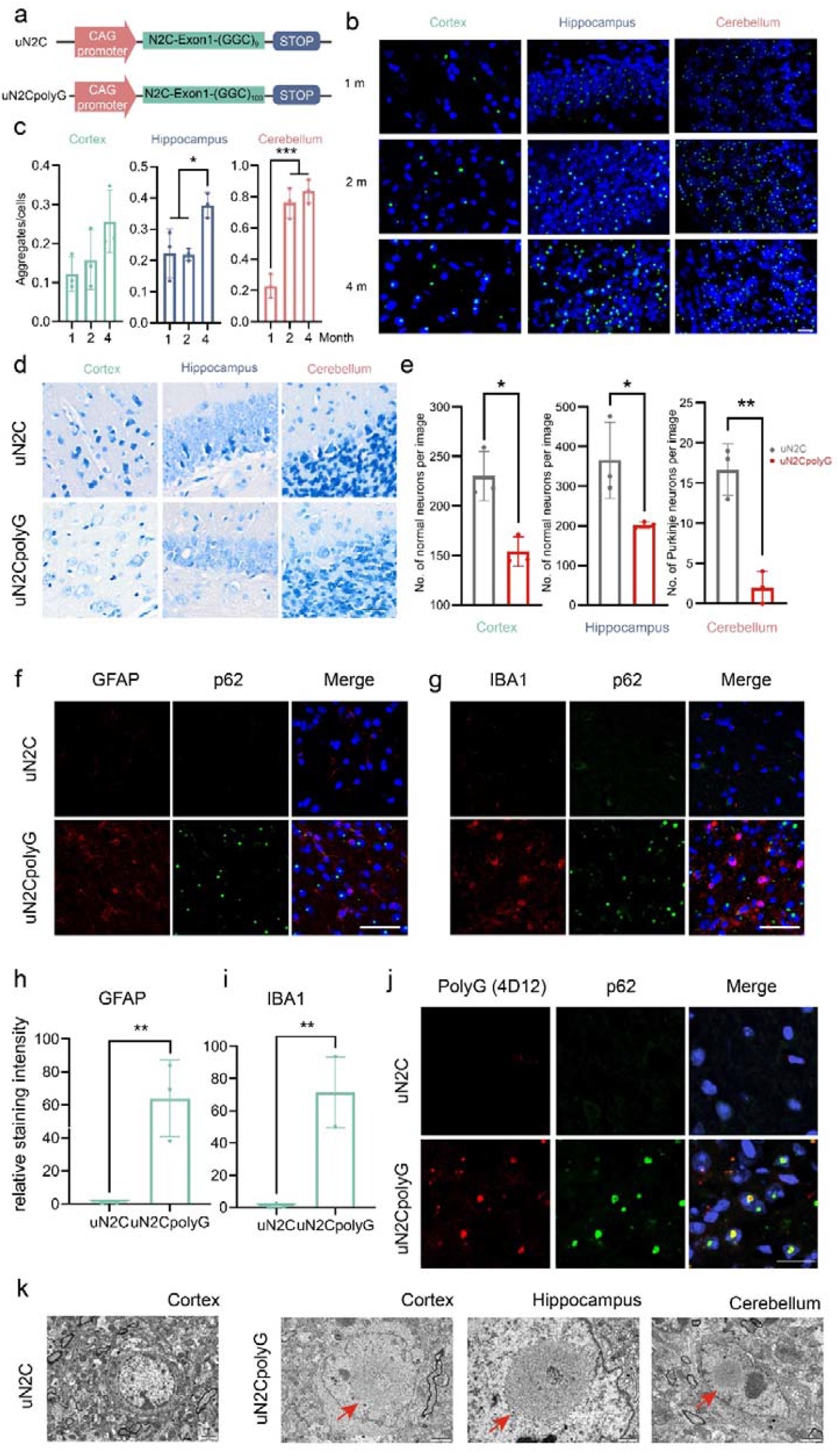
Expression of uN2CpolyG causes progressive neurodegeneration in transgenic mice. (a) Schematic diagram of CAG promoter-uN2C-(CGG)n-stop constructs. Top: Schematic of uN2C. Bottom: Schematic of uN2CpolyG. (b) Immunofluorescent staining against p62 in the different brain regions of uN2CpolyG mice at P30, P60 and P120. Green, p62; blue, DAPI. Scale bar, 20 μm. (c) Quantification of p62 positive inclusions relative to total cells. Data are presented as mean ± SD. [N = 3 mice per group, *P < 0.05 (hippocampus), ***P<0.001 (cerebellum), data was analysed using one-way ANOVA with Dunnett’s multiple comparison]. (d) Representative Nissl staining of the different brain regions of uN2C and uN2CpolyG mice. Scale bar, 100 μm. (e) Cell counts per visual field (×400) found in the slides with Nissl staining on P120. [N = 3 mice per group, *P < 0.05 (cortex and hippocampus), **P<0.01 (cerebellum), data was analysed using unpaired t test]. (f) Representative immunofluorescent staining against GFAP in the cortex of uN2C and uN2CpolyG mice at P120. Red, GFAP; green, p62; blue, DAPI. Scale bar, 50 μm. (g) Representative immunofluorescent staining against IBA1 in the cortex of uN2C and uN2CpolyG mice at P120. Red, IBA1; green, p62; blue, DAPI. Scale bar, 50 μm. (h and i) Quantification of GFAP (h) and Iba1 (i) relative staining intensity. Data are presented as mean ± SD. [N = 3 mice per group, **P < 0.01, data was analysed using unpaired t test]. (j) Immunofluorescence against uN2CpolyG using 4D12 and p62 antibody on brain sections of uN2C and uN2CpolyG mice. Red, uN2CpolyG; green, p62; blue, DAPI. Scale bar, 20μm. (k) Electron microscopy of neurons from the uN2C and uN2CpolyG mice at P120. Red arrows indicate the intranuclear inclusions. Scale bar, 2 μm (cortex and cerebellum) and 1μm (hippocampus).

To determine whether the transgenic mouse model of NIID can faithfully recapitulate the pathological features of neurodegenerative diseases observed in patients [27], we initially performed detailed anatomical examinations on uN2CpolyG mice and their control counterparts, the uN2C mice. The H & E staining showed the typical eosinophilic intranuclear inclusions in cortex, hippocampus, and cerebellum of uN2CpolyG mice (Figure 1c). The immunofluorescence analysis revealed an age-dependent increase in p62-positive inclusions within specific brain regions in uN2CpolyG mice, including the cortex, hippocampus, and cerebellum (Figure 2b, 2c). Compared with uN2C mice, these brain regions exhibited substantial neuronal loss by 4 months in uN2CpolyG mice, as confirmed by Nissl staining (Figure 2d and 2e). Next, we evaluated gliosis, a hallmark of neurodegeneration, by using glial fibrillary acidic protein (GFAP) as an astrocyte marker and ionized calcium-binding adapter molecule 1 (IBA1) as a microglia marker. As anticipated, the activation of both astrocytes and microglia was observed in cortex of uN2CpolyG mice at 4 months of age (Figure 2f, 2g, 2h and 2i). Collectively, these findings suggest that uN2CpolyG expression induces neurodegeneration and neuroinflammation in the nervous system.

Subsequently, we utilized the uN2CpolyG-specific antibody (4D12) [17] to detect the specific expression of polyG in the mouse model. Our findings revealed that polyG colocalizes with the p62 protein in the nucleus (Figure 2j). Furthermore, electron microscopy revealed the presence of filamentous neuronal intranuclear inclusions across multiple brain regions (Figure 2k), indicating that this mouse model successfully recapitulates the pathological phenotype of NIID.

### uN2CpolyG forms intranuclear inclusions in multisystem

NIID is a multisystem disorder characterized by a broad spectrum of clinical manifestations. These include cognitive decline, stroke-like episodes, encephalitic attacks, autonomic dysfunction, muscle weakness, cerebellar ataxia, parkinsonian features, peripheral neuropathy, psychiatric disturbances, visual impairments, and other systemic symptoms [24, 28]. To date, there has been a lack of transgenic mouse models that accurately recapitulate the multisystemic intranuclear inclusions observed in NIID. Here, we aimed to investigate the multisystemic pathology and potential impact of uN2CpolyG expression in a mouse model. Given that muscle involvement and peripheral neuropathy are prominent clinical manifestations in the multisystemic symptoms of NIID, substantially affecting patients’ quality of life, the pathological alterations in both the muscular and nervous systems have become our primary research focus. Histopathological examination and transmission electron microscopy (TEM) revealed preserved muscle architecture in uN2C mouse, demonstrating well-organized muscle fibers without structural abnormalities (Figure 3a-3d). In contrast, H & E staining in the muscle of uN2CpolyG mouse displayed distinct pathological features, including eosinophilic cytoplasmic inclusions and muscle fiber size variation (Figure 3e-3h). Immunofluorescence analysis further confirmed the presence of p62-positive intranuclear aggregates in muscle samples (Figure 3i). According to our previous study, demyelinating degeneration and axonal degeneration could both be observed in NIID patients, even in the absence of clinical neuropathy symptoms [29]. Compared to uN2C mice (Figure 3j-3l), uN2CpolyG mice exhibited mixed demyelination and axonal degeneration, accompanied by p62-positive deposits in the nucleus and filamentous intranuclear inclusions as observed under TEM (Figure 3m-3o). Consistent with the multisystemic pathology observed in NIID patients [28], the p62-positive intranuclear inclusions were frequently detected in heart, liver, lung and kidney tissues (Figure 3p-3t). These results demonstrate that the mouse model comprehensively recapitulates systemic degeneration and the widespread presence of intranuclear inclusions across multiple organs, which are hallmark features of NIID.

**Figure 3.**
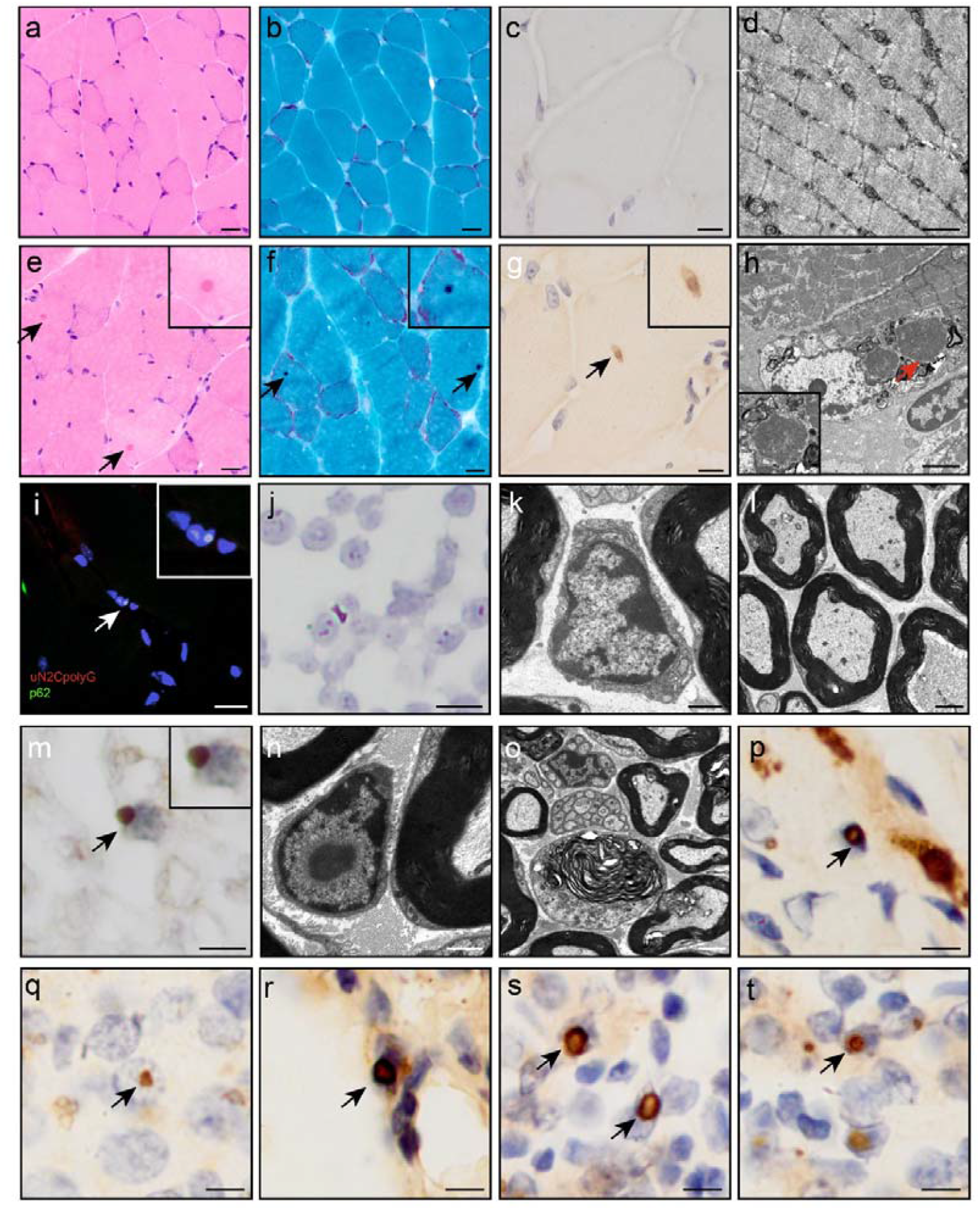
uN2CpolyG forms intranuclear inclusions in multisystem. Systemic pathological analysis of uN2C and uN2CpolyG mice at postnatal day 120 (P120): gastrocnemius muscle sections from uN2C mice with hematoxylin and eosin (H&E) staining (a), modified Gomori trichrome (mGT) staining (b), p62 immunohistochemistry (IHC) (c), and transmission electron microscopy (TEM) (d); uN2CpolyG mice gastrocnemius muscle with H&E (e), mGT (f), p62 IHC (g), TEM (h), and p62 immunofluorescence (IF) (i); sciatic nerve from uN2C mice with p62 IHC (j) and TEM (k,l); uN2CpolyG mice sciatic nerve with p62 IHC (m) and TEM (n,o); and peripheral organ p62 IHC in uN2CpolyG mice including heart (p), liver (q), lung (r), and kidney (s, t). Arrows mark cytoplasmic inclusions in (e-g) and intranuclear inclusions in (h, i, m, p-t). Scale bars: 20 μm (a, b, e, f, i), 10 μm (c, g), 5 μm (j, m, p-s), 2 μm (h, l, o), and 1 μm (d, k, n).

### Expression of uN2CpolyG leads to neurodegenerative phenotypes in transgenic mice

Given that dementia and motor impairments are the primary clinical manifestations in patients with NIID, we aimed to investigate whether our mouse model could replicate these neurological deficits. Our observations revealed that uN2CpolyG transgenic mouse model began to exhibit progressive behavioral abnormalities after 2 months of age, including reduced activity, hunched position and finally lead to cachexia before premature mortality (Figure 4a, 4b). Notably, a significant reduction in body weight was observed in uN2CpolyG mice during the terminal disease phase (Figure 4c). Deterioration of motor performance was observed in the 4-month-old uN2CpolyG mice, which showed decreased grip strength and a shortened latency to fall from a rotarod. These deficits were not detected at 2-month-old uN2CpolyG mice, indicating a progressive deterioration (Figure 4d, 4e). In the open field test, uN2CpolyG mice exhibited a significant reduction in both total activity duration and total distance traveled compared to control mice (Figure 4f-4h), indicating potential declines in motor function. At 4 months of age, uN2CpolyG mice exhibited a significantly reduced correct rate (62.45% ± 10.67) in the Y-maze test compared to control (73.42% ± 6.455) and uN2C groups (73.08% ± 6.648), demonstrating a specific deficit in spatial working memory (Figure 4i-4j).

**Figure 4.**
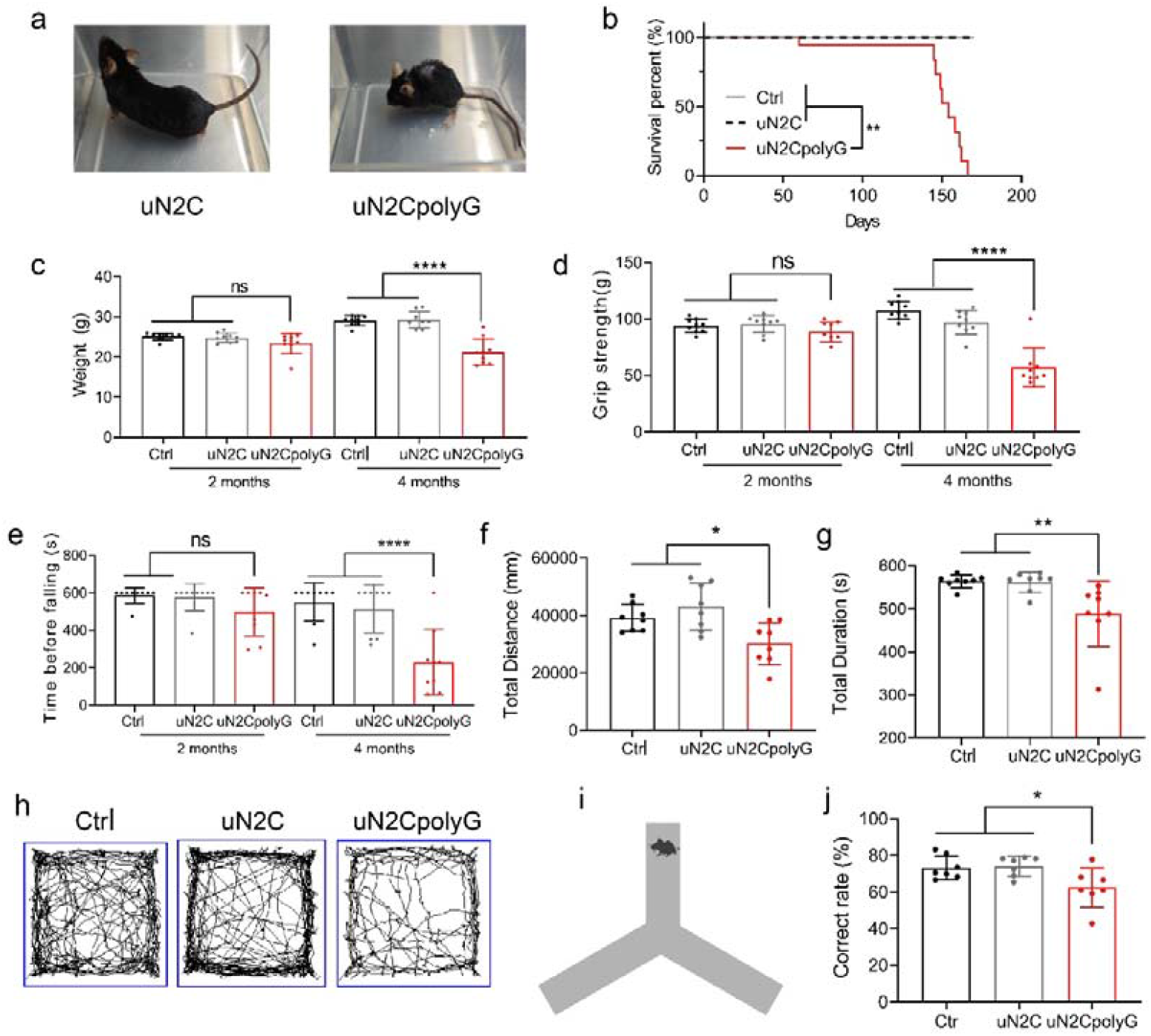
Expression of uN2CpolyG leads to neurodegenerative phenotypes in transgenic mice. (a) Representative image of uN2C and uN2CpolyG mice at P120. (b) Kaplan-Meier survival curve of control, uN2C and uN2CpolyG mice. N=12, **P<0.01. (c-e) Functional assessments at P60 and P120: body weight (c), grip strength (d), and rotarod performance (e). N=9, ****P<0.0001. (f to h) Open field test in control, uN2C and uN2CpolyG mice. Total distance (f) and total duration (g) were measured. N=8, *P<0.05, **P<0.01, ****P<0.0001. (h) Representative image of moving trajectory in the open field test. (i) Schematic diagram of the Y-maze test. (j) Spontaneous alternation behavior in the Y-maze test for control, uN2C, and uN2CpolyG mice. N = 7, *P < 0.05. Data are presented as means ± SD.

## Discussion

NIID is a multisystemic disorder characterized by the presence of ubiquitin- and p62-positive polyglycine intranuclear inclusions [17, 18]. This condition affects not only the nervous system but also various other tissues and organs throughout the body [24, 28]. NIID, as the most prevalent polyG disease in East Asia, serves as an ideal disease model for studying the pathogenesis of this class of diseases. Although the clinical manifestations of polyG diseases are highly heterogeneous, they share a common pathological hallmark: ubiquitin-positive intranuclear inclusions in systemic organs [1, 23]. However, the pathological role of polyG aggregates within the nucleus remains largely elusive. To investigate the multisystemic impairment caused by polyG intranuclear inclusions, we developed a new polyG mouse model. This model aims to enhance our understanding of the mechanisms underlying polyG toxicity across multiple systems. We found that polyG expression leads to the formation of intranuclear inclusions not only in the nervous system but also in various other tissues in this mouse model, thereby recapitulating the polyG pathology observed in NIID patients [1, 28]. The mice exhibited reduced body weight, premature mortality, abundant p62-positive polyG inclusions composed of fibrils, and notable neuronal loss in the cortex, hippocampus, and cerebellum. Additionally, mice expressing uN2CpolyG exhibited behavioral, motor, and memory impairments. The hippocampal neuronal loss observed in these mice likely contributed to their memory deficits, while the depletion of Purkinje cells in the cerebellum—a region classically recognized for its critical role in motor control—may have accounted for their deficits in motor coordination and balance. Neuronal loss was accompanied by the activation of gliosis in mice expressing uN2CpolyG, which is consistent with the observation of upregulated neuroinflammation in NIID [30]. It should be noted that uN2CpolyG aggregates increase in an age-dependent manner, eventually leading to neuronal cell loss and neurodegenerative phenotypes in this mouse model. This further suggests that the polyG toxic protein is the culprit in NIID and serves as a critical therapeutic target for treating this disease.

## Acknowledgement

We appreciated the cooperation of the individuals and their families. We thank Mr. Jin Xu (Peking University First Hospital) for the work in taking electron microscopy pictures, and Ms. Qingqing Wang, Ms. Jing Liu, Ms. Yuehuan Zuo and Ms. Qiurong Zhang (Peking University First Hospital) for their work in preparation for pathological sections. We thank Dr. You Wan, Ms. Feng Tian, Ms. Xin Zheng, Dr. Xianwen Wu, Mr. Kuo Zhang and Ms. Yanping Chen (Peking University Health Science Center) for their guidance on techniques in laboratory animal science and professional care of laboratory animals. We sincerely thank National Human Brain Bank for Development and Function, Chinese Academy of Medical Sciences and Peking Union Medical College, Beijing, China for providing brain tissues.

## Author contributions

J.D., Z.W. and J.Y. contributed to the study conception and design. The molecular experiments were performed and analysed by Y.W., J.D., Z.Y., and J.Y.. Animal experiments were performed and analysis by Y.W., Y. Z., C.G., Y.L., J.W., B.Y. and J.Z.. Bioinformatic analysis was performed by Y.W, F.Z, and J.D.. Figure layout was performed by Y.W., J.D. and J.Y.. Patient recruitment was performed by Z.W.,Y.Y., and D.H.. The first draft of the manuscript was written by J.D., Y.W., J.Y., N.C-B. and all authors commented on previous versions of the manuscript. All authors read and approved the final manuscript.

## Data Availability

The data that support the findings of this study are available from the corresponding author upon reasonable request.

